# Cell-free synthesis and reconstitution of Bax in nanodiscs: comparison between wild-type Bax and a constitutively active mutant

**DOI:** 10.1101/2022.06.09.495475

**Authors:** Akandé Rouchidane Eyitayo, Marie-France Giraud, Laetitia Daury, Olivier Lambert, Cécile Gonzalez, Stéphen Manon

## Abstract

Bax is a major player in the mitochondrial pathway of apoptosis, by permeabilizing the Outer Mitochondrial Membrane (OMM) to various apoptogenic factors, including cytochrome c. In order to get further insight into the structure and function of Bax when it is inserted in the OMM, we attempted to reconstitute Bax in nanodiscs. Cell-free protein synthesis in the presence of nanodiscs did not allow to obtain Bax-containing nanodiscs, but it provided a simple way to purify full-length Bax without any tag. Purified wild-type Bax (BaxWT) and a constitutively active mutant (BaxP168A) displayed biochemical properties that were in line with previous characterizations following their expression in yeast and human cells followed by their reconstitution into liposomes. Both Bax variants were then reconstituted in nanodiscs. Size exclusion chromatography, dynamic light scattering and transmission electron microscopy showed that nanodiscs formed with BaxP168A were larger than nanodiscs formed with BaxWT. This was consistent with the hypothesis that BaxP168A was reconstituted in nanodiscs as an active oligomer.

## Introduction

Bcl-2 family members are major regulators of the intrinsic pathway of apoptosis ([1], for review). Among them, the multidomain pro-apoptotic protein Bax is central to the process [2–4]. Bax is present, under an inactive conformation, in the vast majority of non-apoptotic cells. Following an apoptotic signal, Bax is activated and relocalized at the outer mitochondrial membrane (OMM), where it is oligomerized to form a large size pore [5–8]. The formation of this pore promotes the release of several so-called apoptogenic factors, that contribute to the downstream activation of caspases, and other pro-apoptotic factors.

When in solution, Bax is organized in 9 α-helices [9]. Like for anti-apoptotic proteins Bcl-2 and Bcl-xL, the C-terminal α-helix of Bax, α9, is hydrophobic and has been first considered as an essential membrane anchor. However, contrary to the C-terminal α-helix of Bcl-2 or Bcl-xL, for which the membrane-anchoring function has been supported by clear evidences [10], the actual role of the C-terminal α-helix of Bax as a membrane anchor remains discussed. Early *in vitro* experiments on chimeric proteins between Bax and Bcl-xL showed that the C-terminal α-helix of Bax (BaxHα9) could not serve as an anchor for Bcl-xL, while the C-terminal α-helix of Bcl-xL could serve as a mitochondrial membrane anchor to Bax [11], although the chimera lost its pro-apoptotic properties [12]. The capacity of recombinant BaxΔHα9 (Bax without its C-terminal α-helix) to permeabilize liposomes initially showed conflicting results: Montessuit *et al* found that the protein was 25-fold less active than full-length Bax [13] while Saito *et al* found that the protein was fully active [14]. It was further shown that BaxΔHα9 was indeed inactive but could be activated, through its oligomerization, by the presence of a small amount of octylglucoside [15]. This suggested that BaxHα9 was not involved in Bax insertion but rather in the process of Bax oligomerization. BaxHα9 is very similar to the C-terminal α-helices of both Bcl-2 and Bcl-xL; however, both anti-apoptotic proteins have positively charged residues upstream their C-terminal α-helix that could help stabilizing inserted proteins, contrary to Bax [10]. Beyond this discussion about its role as a *bona fide* membrane anchor, there is ample additional evidence that BaxHα9 plays a role in Bax activation. The residue S184, that is located within BaxHα9, is the target of several protein-kinases, including the survival kinase AKT [16,17]. The phosphorylation of S184, or the substitution of S184 by a negatively charged residue D or E, has been shown to decrease the mitochondrial localization of Bax in human cells [16,18] and following heterologous expression in yeast cells [19]. Besides, Bax phosphorylation correlated with the resistance to BH3-mimetics in primary cancer cells [18] and, conversely, molecules preventing Bax phosphorylation by interacting with the pocket around S184 did favor Bax activation [20]. It should be noted, however, that the consequences of the phosphorylation of S184 on Bax function are still not completely elucidated, because the phosphorylation of S184 may have additional effects on Bax stability, its interaction with anti-apoptotic partners, and its retrotranslocation [21–23].

Another set of site-directed mutagenesis experiments was done on residue P168, that is located in the loop between Hα8 and Hα9. Due to their greater tendency to adopt the *cis* form, proline residues are more subject to support *cis/trans* isomerization, with possible dramatic consequences on protein conformation [24]. In addition, a peptidyl-prolyl *cis/trans* isomerase, Pin1, is able to interact with Bax and modulate its mitochondrial localization [25]. The substitution of P168 by other residues led to conflicting results. When expressed in human glioblastoma, a P168V mutant displayed an increased mitochondrial localization [26]. A similar effect was observed with a P168A mutant expressed in yeast, associated to a dramatic increase of Bax-induced release of cytochrome c [27]. When synthesized as a recombinant His_6_-tagged protein, the same P168A mutant displayed an increased ability (compared to BaxWT) to bind and permeabilize liposomes, yeast mitochondria and human mitochondria (HCT116 Bax^−/−^), although the increase was less marked than in *in cellulo* experiments [28]. It was hypothesized that the isomerization of P168 could significantly modulate the position of Hα9 between a “closed” position, in the hydrophobic groove formed by the three BH domains and an “opened” position, where Hα9 could interact with the lipid bilayer and/or a proteic partner [29]. However, other mutagenesis experiments gave different results. A mutation P168G was shown to stabilize an inactive dimer that remained in the cytosol in MEFs [30]. GFP-BaxP168A or YFP-BaxP168A fusions were also found to remain cytosolic in staurosporin-treated MEFs [31], in mouse FSK7 cells [32] and in human HEK293 cells [33]. It is possible that still undefined other factors, beyond the conformation of Bax itself, are involved in the localization of this mutant. Alternatively, the presence of additional sequences at the N-terminal end (GFP, His_6_ tag) or the complete deletion of the mobile N-terminal sequence may modulate conformational changes of the C-terminus: indeed, it has been shown that the activation of Bax could be modulated by an interaction between R9 and D154 [19] and that Bax was strongly activated by the deletion of its 20 N-terminal residues [33].

The structure of full-length soluble monomeric Bax (supposedly inactive) has been solved by NMR and showed the localization of Hα9 in the hydrophobic groove (the “closed” position defined above) [9]. The structure of a Bax dimer obtained after its activation by a peptide covering the BH3 domain of Bid has been solved by X-Ray cristallography, but this structure lacked the Hα9 [34]. The structure of the inactive dimer containing the P168G mutation has also been solved by X-Ray cristallography, showing the Hα9 in the hydrophobic groove [30].

The structure of the full-length and active Bax (forming the pore) is then still missing. Biochemistry experiments [15] suggest that it is an oligomer (260kDa) containing about 12 Bax monomers (21kDa). Electrophysiology experiments showed that Bax could form a pore having a size of about 4.6 nm in diameter [6,35], large enough to permeabilize the membrane to a small protein like cytochrome c (12.5kDa, ~3nm). It is noteworthy that the degree of oligomerization may increase with time [36]. The structures obtained by high resolution fluorescence microscopy and atomic force microscopy showed much larger structures, with diameters from 40 to more than 100nm [7,8], that are unlikely to be formed during the early steps of apoptosis, when there is a selectivity of released proteins: for example, mature smac/diablo (a 21kDa protein) is released 4 times more slowly than cytochrome c (12.5kDa) [37]. Even more strikingly, when expressed in yeast, Bax is able to release cytochrome c but not a fusion cytochrome c-GFP [38]. These observations are thus not compatible with the existence of an unselective pore having a diameter larger than 5nm.

It is therefore desirable to set up a controlled model of active Bax oligomers, both for biochemical and structural studies. In recent years, nanodiscs have been developped as a powerful model to study membrane proteins [39]. The diameter of nanodiscs is controlled namely by the size of the scaffold protein [40]. Their stability and relative homogeneity has allowed investigators to get both biochemical and structural data on a large number of membrane proteins [41].

In the present study, we first attempted to use cell-free protein synthesis in the presence of pre-formed nanodiscs to combine the translation and the insertion of Bax into nanodiscs, as it had been done for other membrane proteins [42–44]. This did not work as intended initially, as we observed that the presence of nanodiscs induced a precipitation of Bax. The fact that the precipitation occurred at lower nanodisc concentration for BaxWT than for the mutant BaxP168A suggested that this precipitation was linked to Bax propensity to form oligomers that could not be inserted. Interestingly, nanodisc-induced Bax precipitation could be fully prevented by the co-synthesis with its anti-apoptotic partner Bcl-xL.

In these experiments, Bax precipitation provided the opportunity to quickly produce purified untagged full-length BaxWT and the constitutively active BaxP168A mutant. Like it was previously shown for the His_6_-tagged proteins produced in the presence of low amounts of Brij-58 or F8-TAC [28], the BaxP168A mutant was more active than BaxWT, showing that the difference observed *in cellulo* was conserved on recombinant proteins. Both proteins were further used to co-form Bax-containing nanodiscs. The comparison of the size of nanodiscs formed with BaxWT or with the mutant BaxP168A suggested that the later contained oligomeric active Bax, consistent with previously published electrophysiology and biochemistry experiments on mitochondria-inserted Bax.

## Results

### Effect of the presence of pre-formed nanodiscs on Bax cell-free synthesis

Our initial goal was to produce untagged full-length Bax in a cell-free protein synthesis system, in the presence of pre-formed nanodiscs, in order to get Bax as a nanodisc-inserted fraction. The rationale was that the neo-synthesized peptide chain would be simultaneously integrated in the lipid bilayer of nanodiscs, so that hydrophobic domains were not exposed to the aqueous environment, thus preventing mis-folding and aggregation. We used a Bax mutant, carrying a single substitution P168A, that was previously shown to be spontaneously inserted in the mitochondrial outer membrane when expressed in mammalian cells [26] and yeast cells [27]. A His_6_-tagged version of this mutant, produced in a cell-free system in the presence of a small amount of the detergent Brij-58 or the polyfluorinated surfactant F8-TAC, was shown to permeabilize liposomes and isolated mammalian and yeast mitochondria [28].

After a 20 hours production, the reaction mix was centrifuged, and the pellet and supernatant were analyzed by Coomassie Blue staining (Fig.1A). Unexpectedly, we observed that the presence of nanodiscs led to an increased proportion of BaxP168A in the pellet.

**Figure 1:**
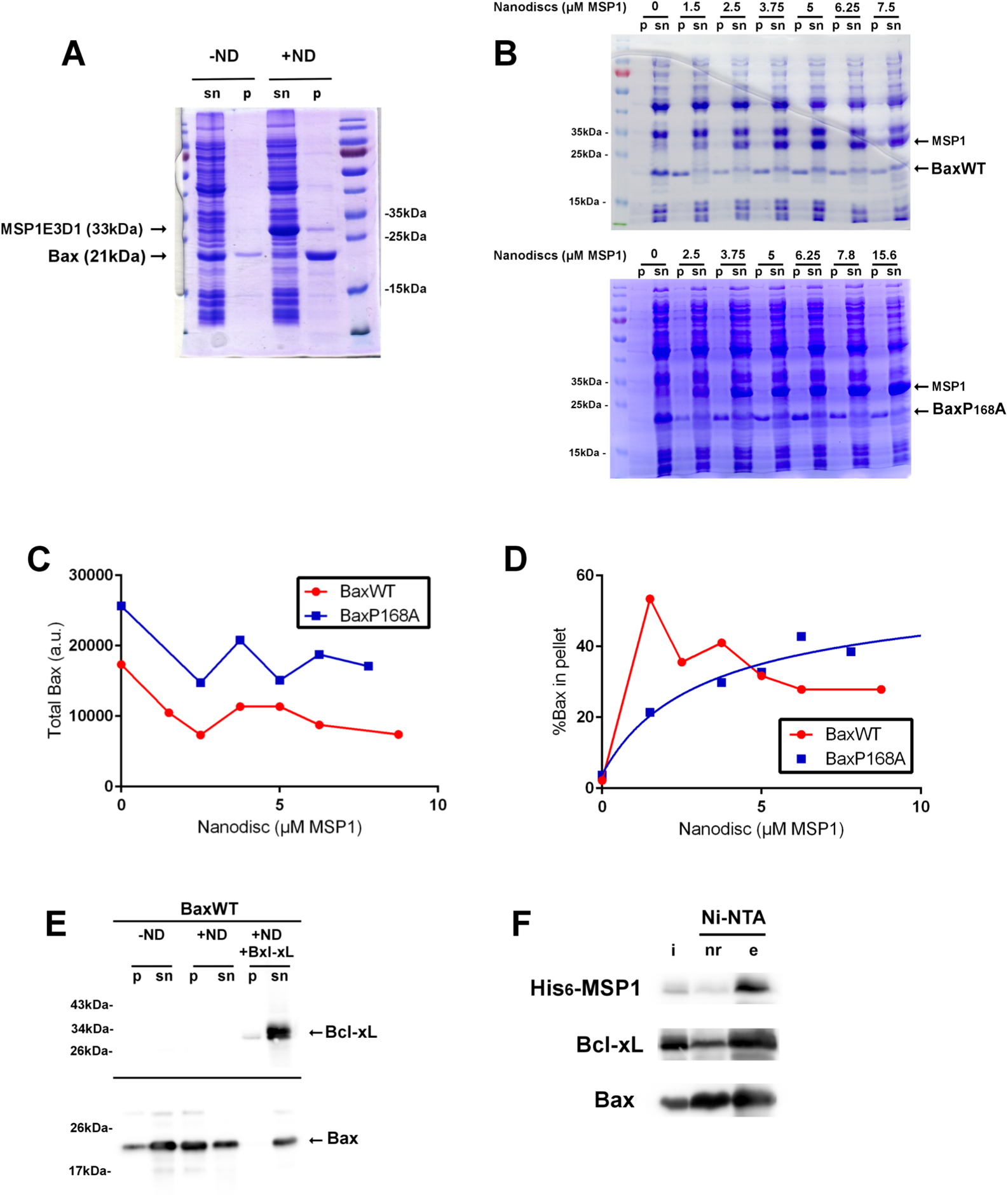
The presence of nanodiscs induces the precipitation of Bax during cell-free protein synthesis. (A) BaxP168A was synthesized in the absence (-ND) or in the presence (+ND) of nanodiscs. The reaction mix was centrifuged (20,000 x g; 10 minutes) and the supernatant (sn) and pellet (p) were solubilized in Laemmli buffer, ran on SDS-PAGE and stained with Coomassie Blue. (B) Cell-free synthesis of BaxWT (top) or BaxP168A (bottom) was done in the presence of a range of nanodisc concentrations (expressed as µM MSP1). After the synthesis, reaction mixes were separated as above and stained with Coomassie Blue. The same proportions of resuspended pellets and supernatants were loaded. (C,D) The amounts of Bax in the pellets and supernatants were measured by the densitometry of Coomassie Blue-stained gels, and plotted v/s the concentration of nanodiscs. (C) Total Bax = Bax in pellet+Bax in supernatant. (D) Bax in pellet/Total Bax. (E,F) BaxWT and Bcl-xL were co-synthesized in a cell-free assay. Pellets and supernatants were analyzed by western-blots. The supernatants containing nanodiscs (i) were pulled-down with Ni-NTA-sepharose and the presence of His_6_-MSP1, Bcl-xL and BaxWT was measured in the non-retained fraction (nr) and the eluate (e).

We next tested if this behavior was due to the peculiar property of BaxP168A, that is spontaneously more membrane-associated than wild-type Bax. A range of nanodiscs concentration was added to test their ability to induce the precipitation of both BaxWT and BaxP168A (Fig.1B-D). We first observed that, in the presence of nanodiscs, the total amount of synthesized Bax (supernatant+pellet) tended to decrease (Fig.1B,C). The second observation is that, like BaxP168A, BaxWT precipitated in the presence of nanodiscs (Fig.1B,D). However, while the proportion of precipitated BaxP168A increased with the concentration of nanodiscs, the precipitation of BaxWT was maximal at a low nanodiscs concentration (1.5µM MSP1) and then decreased (Fig.1D). Note that the range of nanodisc concentrations inducing these effects may slightly vary between experiments. This was likely due to the “quality” of nanodiscs preparations. However, the general behavior and the difference between BaxWT and BaxP168A was highly reproducible.

We hypothesized that, in the presence of the lipid bilayer provided by a small amount of nanodiscs, BaxWT acquired a conformation that limited its solubility, but remained poorly able to be inserted into nanodiscs, thus promoting its precipitation. The effect was less dramatic with BaxP168A, which can be, at least partly, inserted into nanodiscs. To support this hypothesis, we did co-synthesis experiments with BaxWT and Bcl-xL. Indeed, we have previously shown that Bcl-xL stimulated the localization of BaxWT in the outer mitochondrial membrane of non-apoptotic mammalian cells and of yeast cells [45]. When both proteins were synthesized together in the presence of nanodiscs, BaxWT was not precipitated and both proteins were found in the supernatant (Fig.1E). Furthermore, pull-down against His_6_-MSP1 showed that the vast majority of Bcl-xL and a large proportion of BaxWT were indeed associated with nanodiscs (Fig.1F).

### Resolubilisation of precipitated Bax by detergents

When synthesized in the presence of nanodiscs, pelleted Bax was largely devoid of contaminants from the bacterial lysate (Fig.1A). This purity could be further improved by quickly washing the pellet in the presence of 500mM NaCl (Fig.2A,B). The high degree of purity opened an opportunity to recover purified full-length Bax without any additional residue. We thus attempted to resolubilize it to test if the protein could be used for further biochemical studies. Both 0.1% Nonidet P40 (NP-40) (Fig.2C) and 0.05% Brij-58 (Fig.2D) efficiently resolubilized both BaxWT and BaxP168A. Since we have previously shown that Brij-58 adequatly maintained the properties of His_6_-tagged Bax, and namely the distinct properties of BaxWT and BaxP168A [28], we selected this detergent for further studies on resolubilized untagged Bax.

**Figure 2:**
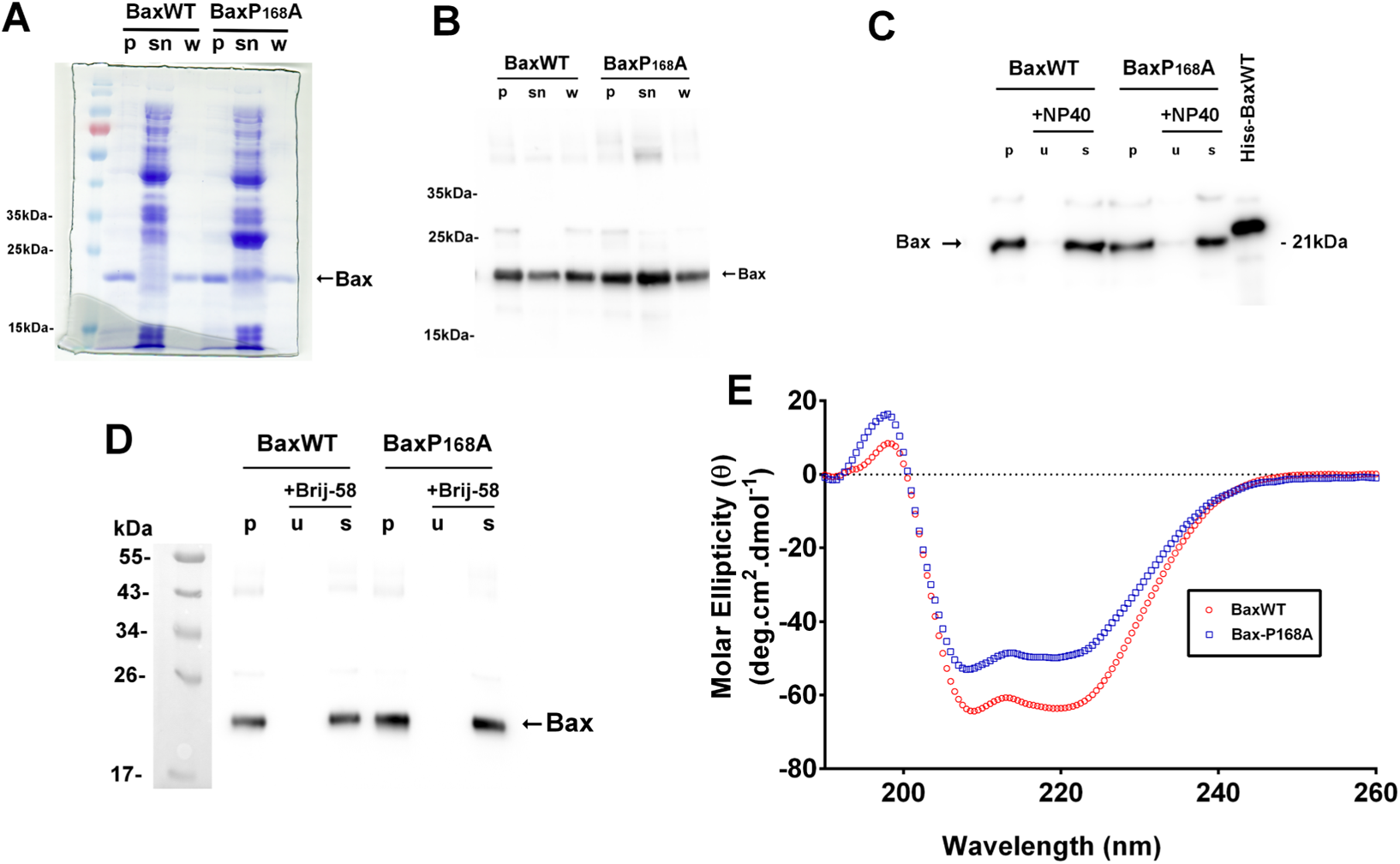
Resolubilization of precipitated Bax. (A,B) After the cell-free synthesis, the pellet (p) and supernatant (sn) were separated as in Figure 1. The pellet was quickly washed with 500mM NaCl and re-centrifuged (w). After separation on SDS-PAGE, proteins were revealed by Coomassie Blue staining (A) or Western-Blotting against Bax (B). (C,D) Washed pellets (w) were added with a 10mM Na/Hepes buffer (pH 7.6) containing 250mM NaCl, 1mM EDTA and 0.05% Nonidet-P40 (C) or 0.05% Brij-58 (D). The solubilization was done overnight at 4°C under gentle agitation. After a 20,000 x g, 10 min centrifugation, the pellet of unsolubilized material (u) and solubilized proteins (s) were separated on SDS-PAGE and revealed by a Western-Blot against Bax. The lane labeled “His_6_-BaxWT” contained a known amount of the His_6_-tagged protein produced in the presence of F8-TAC and purified on His-Trap, and served as a quantification standard. (E) Resolubilized proteins were dialyzed twice against a 40mM Sodium Phosphate buffer (pH 7.2). The protein concentration were 84 and 67µg/mL for BaxWT and BaxP168A, respectively. Spectra were acquired between 260 and 190nm at room temperature (20°C) and corrected with the baseline.

The conformation of resolubilized Bax was assessed by CD spectroscopy (Fig.2E). Resolubilized Bax was first dialyzed against 40mM sodium phosphate (pH 7.2), to eliminate Cl^−^ ions. Both solubilized proteins generated CD spectra that were compatible with a large proportion of α-helices (~70%) (Table 1). These values were similar to those previously acquired with His_6_-Bax produced in cell-free synthesis in the presence of 0.01% Brij-58 or 0.05% F8-TAC [28] and are consistent with the known Bax structures [9,34]. From these results, Bax precipitated and resolubilized did not significantly differ, in terms of CD spectra, from His_6_-Bax that has been maintained in solution along the purification process.

The next step was to test if resolubilized Bax kept its functional properties. Bax activity was assayed as its capacity to permeabilize liposomes loaded with FITC-Dextran 10kDa, which size is close to that of cytochrome c (12.5kDa). Both BaxWT and BaxP168A induced a permeabilization of liposomes, with a slightly greater effect of the mutant (Fig.3AB). This modest difference between BaxWT and BaxP168A had already been observed for His_6_-tagged proteins synthesized in the presence of Brij-58 (or F8-TAC) and tested on liposomes loaded with small molecules ANTS/DPX (<1kDa) [28].

**Figure 3:**
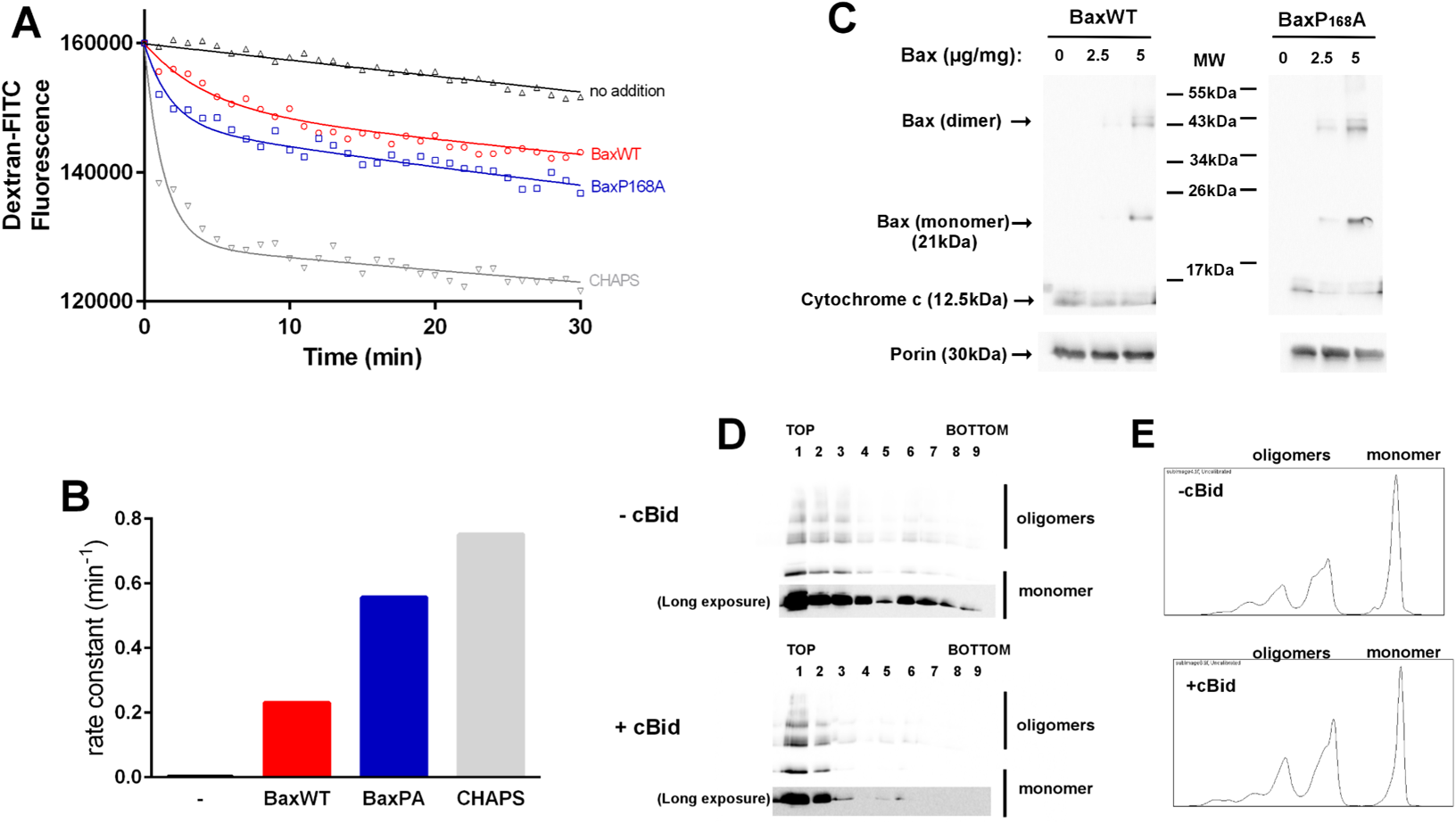
Properties of resolubilized BaxWT and BaxP168A. (A,B) Liposomes loaded with 10kDa Dextran-FITC were prepared as indicated in the Methods section. A mixture containing loaded LUV and an anti-FITC antibody was prepared and displayed in a 96-well black microplate. LUV buffer was added to adjust the final volume at 100µL. At time 0, 100ng BaxWT or BaxP168A or 0.05% CHAPS were added and the fluorescence was measured in a fluorescence plate reader (ex: 495nm; em: 521nm) over time. Curves were fitted with Grafpad software and the initial rates were calculated (B), assuming a one-phase (no addition) or a two-phases decay. (C) The permeabilization of yeast mitochondria was done as described in the Methods section. The pellets were ran on SDS-PAGE and Bax, cytochrome c and Porin (as a control) were revealed by Western-Blots. (D) BaxWT incubated in the presence of LUV, in the absence (-cBid) or in the presence (+cBid) of caspase-3-cleaved Bid (molar ratio Bax/cBid=5). After the incubation, LUV were mixed with 80% Histodenz in LUV buffer and layered at the bottom of ultracentrifuge tubes. The mix was overlayed with 30% Histodenz in LUV buffer and with LUV buffer alone. After the ultracentrifugation, the gradients were recovered as 9 fractions, proteins were precipitated, separated on SDS-PAGE and Bax was revealed by Western Blot. Where indicated, the blotting membranes were overexposed to highlight the presence of Bax in more dense fractions. (E) Normally exposed lanes 1 of each membrane in (D) were analysed by densitometry. The presence of cBid increased the proportion of oligomers by 20-30%.

The capacity of resolubilized Bax to bind and permeabilize isolated mitochondria was next tested. For practical reasons, we used isolated yeast mitochondria, that behave like mammalian mitochondria in *in vitro* assays [28]. Both BaxWT and BaxP168A bound to mitochondria and induced the release of cytochrome c (Fig.3C) with, as expected, a slightly greater efficiency of the mutant BaxP168A. Flotation assays were done to further analyze the capacity of resolubilized Bax to associate with liposomes. A mixture of liposomes and Bax was layered at the bottom of a Histodenz gradient. After centrifugation, Bax was mostly found in the top fractions (Fig.3D), showing that it was associated with liposomes. As expected, the association was significantly stimulated by the presence of cBid. Also, cBid slightly increased the proportion of dimeric/oligomeric forms of Bax (Fig.3E).

From these data, resolubilized BaxWT and BaxP168A both retained similar biochemical properties as His_6_-tagged proteins that were previously produced in cell-free protein synthesis in the absence of precipitation [28], suggesting that the precipitation/resolubilization cycle did not significantly alter BaxWT and BaxP168A properties, and that these proteins could be used in reconstitution assays.

### Co-formation of Bax-containing nanodiscs

The excess of Brij-58 was first eliminated by a single passage on detergent removal columns. 50 to 60% of the proteins were recovered after this step, after which the proteins were used immediately, to avoid further precipitation. Nanodiscs of POPC/POPG 3:1 (w/w) and MSP1E3D1 were then co-formed in the presence of BaxWT or BaxP168A by removing cholate with Bio-Beads. Assays have also been done to remove detergents by overnight dialysis, and gave essentially similar results in the following experiments. After centrifugation (20,000 x g, 15 min) and filtration (0.2 µm), the mixture was loaded on a Sephadex G200 10/300GL column. Typical SEC profiles are shown in Fig.4A.

**Figure 4:**
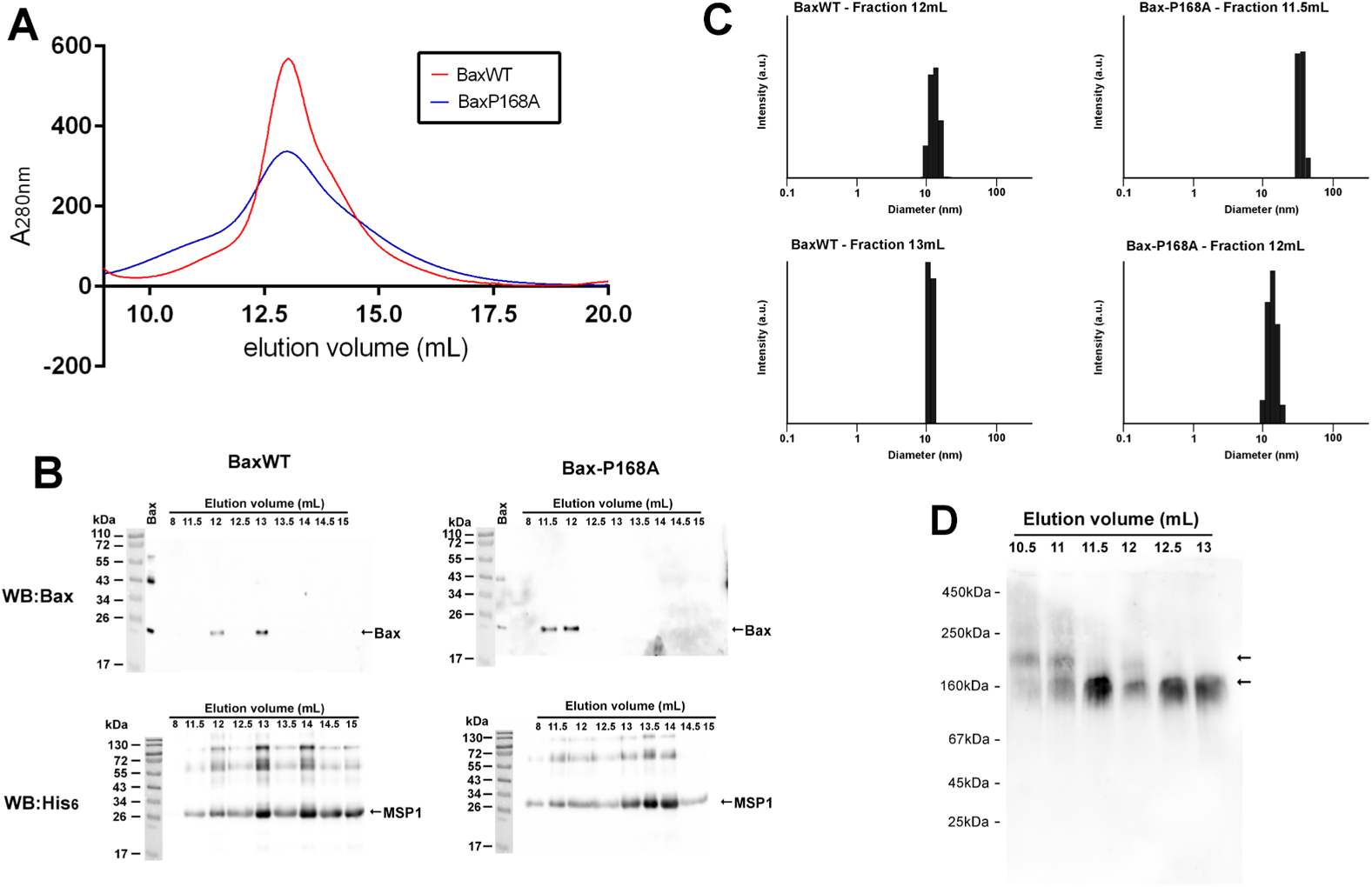
Co-formation of Bax-containing nanodiscs. (A) Bax and nanodiscs were co-formed as described in the method section. Nanodiscs were then loaded on a SEC column (Superdex 200 10/300GL, GE Healthcare) connected to an Äkta purifier. Fractions were eluted with a 10mM Tris/HCl, pH 8.0 buffer containing 100mM NaCl, at a constant rate of 0.5mL/min. Selected fractions were analyzed by Western Blot against Bax and His_6_-MSP1 (B) and by DLS (C). In (B), the lane labeled “Bax” contains the resolubilized protein not reconstituted in nanodiscs. (D) CN-PAGE of SEC fractions of empty nanodiscs were blotted and revealed with anti-His6 antibody, showing 2 populations.

A first peak around 6-8mL contained objects with a size well above 100nm (measured by DLS, not shown) and likely corresponded to MLV that do not contain any detectable amount of Bax (Fraction 8mL in Fig.4B). Main fractions were eluted between 10mL and 14mL with a shoulder between 10mL and 12mL that was higher for BaxP168A, compared to BaxWT.

Fractions from BaxWT reconstitution eluted between 11mL and 14 mL contained objects having an apparent size close to the expected size of nanodiscs, with an apparent diameter close to 12.5 nm (Fig.4C). Western-blot analysis of these fractions showed that the earlier fractions of the peak (12mL and 13mL) contained both Bax and MSP1 while later fraction (13.5mL to 15mL) contained only MSP1. This suggested that a separation occurred between slightly larger nanodiscs containing Bax and smaller empty nanodiscs. The presence of two distinct populations of BaxWT-containing nanodiscs (fractions 12mL and 13mL) was not related to the presence of Bax: indeed, two distinct populations of empty nanodiscs (without Bax) were also observed by CN-PAGE (Fig.4D).

For BaxP168A, an additional early fraction appeared (Fraction 12.5mL) that contained larger objects with an apparent average diameter above 20 nm (Fig.4C). Fraction 12mL contained objects having the same size as Fraction 12mL of BaxWT, around 12.5 nm. Both fractions 11.5mL and 12mL contained Bax and MSP1 (Fig.4B) while later fractions, like for BaxWT, contained only MSP1.

Fractions containing both Bax and MSP1 were then observed by TEM after negative staining. Both fractions from BaxWT and BaxP168A contained objects having the appearance and expected size of nanodiscs (Fig.5A,B). The area of objects was measured and their average diameter was calculated (considering them as disks) (Fig.5C). As expected from the western-blots of SEC fractions, the size of BaxP168A nanodiscs was larger than the size of BaxWT nanodiscs. This is consistent with the hypothesis that BaxP168A tends to form stable oligomers that might generate physical constraints leading to significantly larger nanodiscs. This is in accordance with the biochemical properties of BaxP168A, that is spontaneously more active than BaxWT, as reported above (Fig.3) and in previous studies [28]. Furthermore, a set of co-formation experiments has been done with His_6_-tagged Bax produced in the presence of StrepII-tagged MSP1 (see material and methods). Although most of Bax was precipitated (like in the experiments with untagged Bax and His_6_-MSP1), the small amount of Bax-containing nanodiscs present in the supernatant was next purified by two successive affinity chromatographies on Strep-Tactin and Ni-NTA, and observed by TEM. The very low yield of these experiments only permitted a limited number of observations. Nevertheless, the same difference of size was observed between nanodiscs containing BaxWT and BaxP168A (Fig.5C *v/s* Fig.5D).

**Figure 5:**
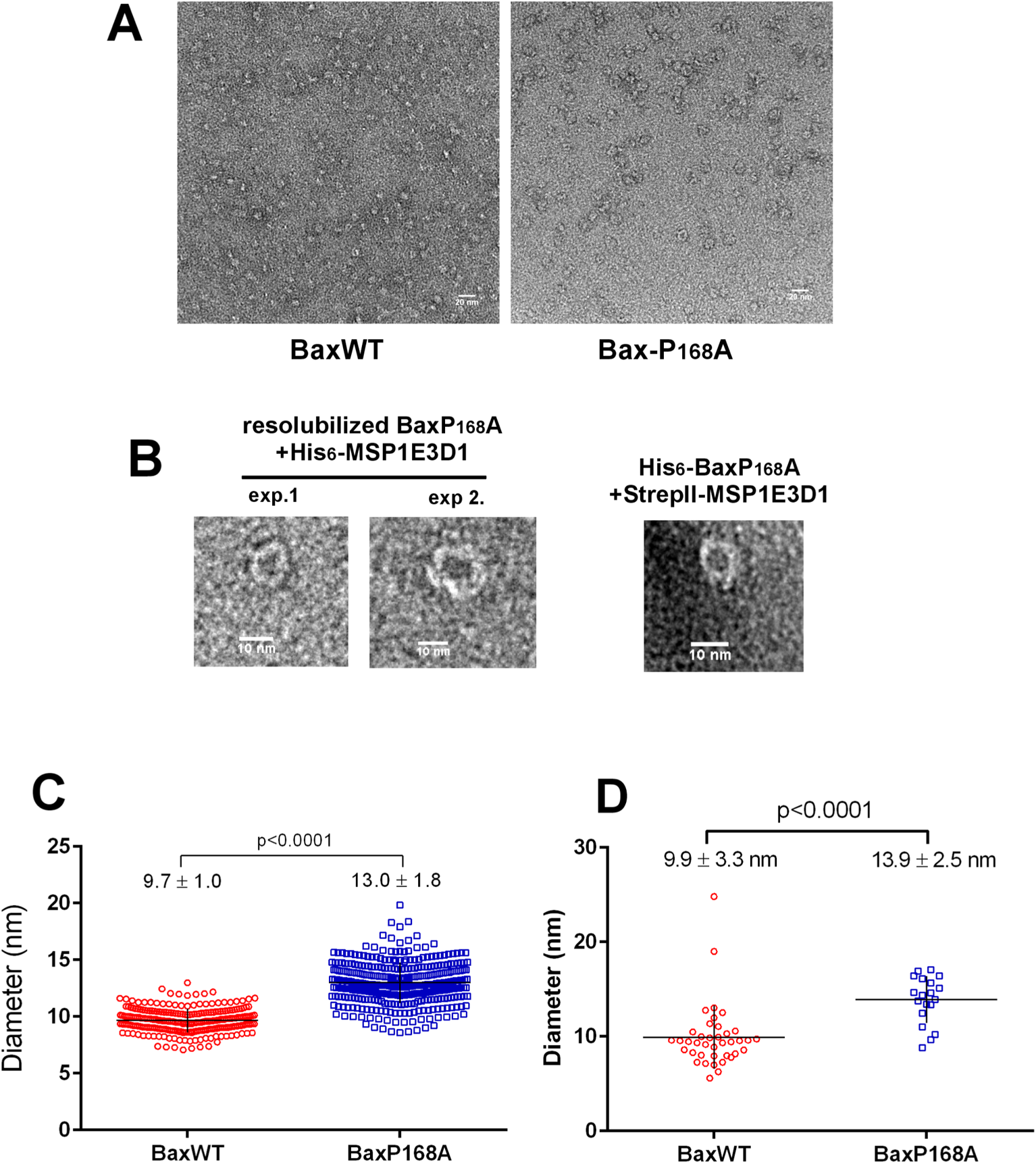
TEM of Bax-containing nanodiscs. (A) TEM images of fractions 12mL (see Figure 4) of BaxWT and BaxP168A. (B) Magnification of individual BaxP168A-containing nanodiscs obtained with precipitated and resolubilized untagged BaxP168A in His_6_-MSPE3D1 nanodiscs purified by SEC in two independent experiments, or with His6-BaxP168A in StrepII-MSP1E3D1 nanodiscs doubly purified by Strep-Tactin and Ni-NTA affinity chromatographies (see methods). (C) Size quantification of nanodiscs was done manually on dm3 TEM images. The perimeter of nanodiscs was drawn and the measured surface was converted to diameters by assuming that they were perfect discs. Both distributions passed the D’Agostino’s K-squared normality test. (D) Size quantification of nanodiscs was done manually on dm3 TEM images. The perimeter of nanodiscs was drawn and the measured surface was converted to diameters by assuming that they were perfect discs. The distribution of BaxP168A nanodiscs passed the D’Agostino’s K-squared normality test.

### Functional Bax properties in nanodiscs

The BaxP168A mutant was first characterized by its higher spontaneous mitochondrial localization and its greater capacity to permeabilize the outer mitochondrial membrane when expressed in yeast [27] or mammalian cells [26]. In a previous study, we observed that the His_6_-tagged version of BaxP168A produced in cell free exhibited an increased recognition by the 6A7 monoclonal antibody [28], that has a greater affinity for active Bax [46]. We then tested if the same property was observed when the protein was reconstituted in nanodiscs (Fig.6A). The ratio of Bax immunoprecipitated by the 6A7 antibody over the ratio of Bax immunoprecipitated by the 2D2 antibody was higher for BaxP168A than for BaxWT (Fig.6B). Of note, the difference between the two proteins was very similar to that previously observed for the His_6_-tagged proteins in solution in the presence of Brij-58 [28] (Fig.6B). The anti-apoptotic protein Bcl-xL is one of the main partner of Bax, that prevents the formation of the large-size pore initiating the release of apoptogenic factors. The mechanism of action of Bcl-xL on Bax is still discussed: the proteins are able to interact in solution (*e.g.* [47]), Bcl-xL stimulates the retrotranslocation of Bax from the OMM to the cytosol [48], but it has also been shown that Bcl-xL could stimulate the mitochondrial translocation of Bax in non-apoptotic cells [45], thus promoting the engagement of Bax in an inactive membrane-associated heterodimer, from which active Bax can be further released by a BH3 peptide or a BH3-mimetic [49].

**Figure 6:**
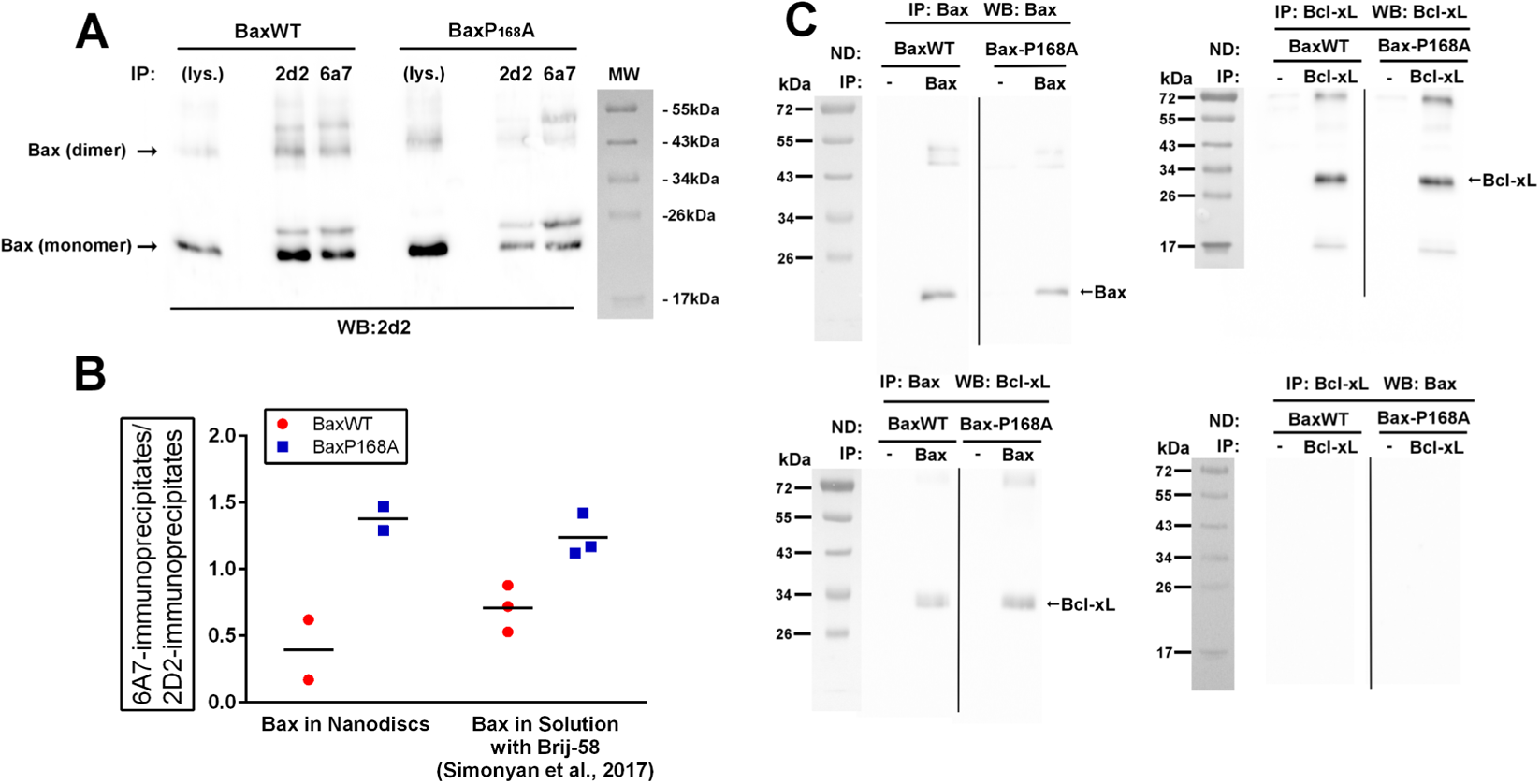
Functional properties of Bax in nanodiscs. (A,B) BaxWT or BaxP168A-containing nanodiscs were incubated overnight with mouse monoclonal Anti-Bax antibodies 6A7 or 2D2, that recognize active Bax or total Bax, respectively [54]. Immunoprecipitates were loaded on SDS-PAGE and revealed by Western-Blot with the 2D2 antibody. (B) The ratio 6A7/2D2 was measured for both BaxWT and BaxP168A, and was compared with previous experiments done with His_6_-tagged proteins produced in the presence of Brij-58 [28]. (C) BaxWT or BaxP168A-containing nanodiscs were incubated overnight in the absence or in the presence of anti-Bax (2D2) or anti-Bcl-xL (E18) antibodies. The immunoprecipitates were loaded on SDS-PAGE and revealed by Western-Blots against Bax or Bcl-xL.

BaxWT and BaxP168A nanodiscs were incubated in the presence of recombinant Bcl-xL, and the co-immunoprecipitation of the two proteins was measured (Fig.6C). Antibodies directed against Bax allowed the co-immunoprecipitation of Bcl-xL, suggesting that Bcl-xL was able to interact with Bax reconstituted in nanodiscs. There was no obvious difference between BaxWT and BaxP168A. Conversely, we could not depict Bax in Bcl-xL-immunoprecipitates, even after an extended exposure. These results suggested that the two proteins were still able to interact but that the conformation of Bax in the nanodisc bilayer may somehow decrease the affinity between Bax and Bcl-xL, and jeopardize the co-immunoprecipitation of Bax by the Bcl-xL antibody.

## Discussion

In a previous study, we showed that Bax could be produced in a cell-free protein synthesis system as a soluble protein, in the presence of a small amount of the detergent Brij-58 or the fluorinated surfactant F8-TAC [28]. The protein was produced as a His_6_-tagged protein to facilitate further purification from the bacterial lysate. The comparison between the BaxWT protein and the constitutively more active mutant BaxP168A showed that the native properties of the protein were conserved. The initial goal of the present study was to associate the cell-free protein synthesis system and the presence of nanodiscs to reconstitute the protein at the same time as it was translated, as it has been done for other proteins [42–44]. Rather unexpectedly, we observed that the presence of nanodiscs during the synthesis induced the precipitation of Bax. By modulating the concentration of nanodiscs, we observed that the “optimal” concentration (*i.e.* inducing the maximal precipitation) was different for BaxWT and BaxP168A (Fig.1D): it was about 1.5µM (measured as MSP1 concentration) for BaxWT, while a plateau above 8µM was observed for BaxP168A (higher concentrations of nanodiscs could not be easily tested, because the cell free assay only permitted a small volume of “custom” additions). The fact that a small addition of nanodiscs is sufficient to initiate a high precipitation ratio of BaxWT might be related to the presence of the lipid bilayer, that may initiate a partial conformational change of Bax which, however does not go all the way until the fully inserted protein, because of the absence of contributing factors, such as the mitochondrial receptor Tom22 [50–52]. In accordance to this hypothesis, the precipitation occurs at higher nanodiscs concentrations for BaxP168A that is able to insert spontaneously in lipid bilayers. The precipitation of BaxP168A may thus occur only when nanodiscs are saturated, that would explain the different shapes of the curves for the two Bax variants. Interestingly, it has been previously shown on isolated mitochondria, that once a fraction of Bax is membrane-inserted, it can serve as a receptor for soluble Bax [53]. This might explain why, after a peak at low MSP1 concentrations, BaxWT precipitation decreased at higher concentration and then stabilized at a level close to that of BaxP168A: this might correspond to the insertion of enough Bax molecules, serving as receptors to the next neosynthesized ones. In the case of P168A, that has spontaneously the adequate conformation for its insertion, the precipitation curve increased continuously. This hypothesis was supported by the fact that the co-synthesis of BaxWT with Bcl-xL in the presence of nanodiscs prevented the precipitation of BaxWT. We have reported previously that, both in non-apoptotic mammalian cells and in yeast cells, the co-expression of Bcl-xL increased the localization of BaxWT in the outer mitochondrial membrane [45]. The fact that this effect occurred in yeast cells, that do not express any Bcl-2 family member, suggested that it depended only on the two proteins. This is confirmed by these cell-free synthesis assays in the presence of nanodiscs, where no additionnal partner of BaxWT and Bcl-xL are present. Since Bax was highly purified in the pellet, we used this opportunity to produce full-length Bax without any tag. As described previously [28], Bax can be produced in cell-free synthesis as a N-terminal His_6_-tagged protein with a high yield and a high degree of purity. His_6_-tagged Bax keeps its functionnal properties (permeabilization of liposomes and mitochondria, interaction with Bcl-xL) and the expected differences between His_6_-BaxWT and His_6_-BaxP168A were observed, although somewhat attenuated as compared to the proteins expressed in yeast or human cells. An important regulation of Bax activation is linked to the movement of the unstructured N-terminal part. Indeed, it has been shown that the deletion of the 20 N-terminal residues of Bax generated a protein having a much greater capacity to bind to mitochondria and to permeabilize the OMM in both yeast and human cells [33,54] and to induce apoptosis in human cells [33,55]. This domain has been named ART (Apoptotic Regulation of Targeting) [54]. Interestingly, a similar splicing variant (called BaxΨ) was found in low-grade glioblastomas showing a greater rate of apoptosis [56]. This higher activity of N-truncated Bax might be related to the observation that the R9 residue, located in the ART, might interact with the D154 residue located in the loop between helices α7 and α8: indeed, the inversion of one of these charges (E9 or K154) induced the activation of Bax, while a double inversion (E9/K154) had the same activity as the wild-type protein (R9/D154) [19]. This suggests that the presence of a positively charged His_6_-tag at the N-terminus might potentially interfere with Bax activation. It is thus desirable to produce the untagged protein. The introduction of a cleavable tag (with a Factor X-recognition sequence) has been done, but the low efficacy of the cleavage led to a 50% decrease of the yield of the protein (Simonyan and Manon, unpublished data) and two additional residues remain after the cleavage. The method of choice to produce untagged Bax has been the fusion with intein, allowing the purification on chitin-sepharose beads and the autocleavage of the tag in the presence of reducers β-mercaptoethanol or DTT [9]. However, the method is not without inconvenients: the production yield is low, the efficacy of cleavage is lower than 50% and, due to the low yield, additional purification steps are required to eliminate several bacterial proteins.

Because we had previously shown that Brij-58 preserved Bax functions [28], this detergent was used to resolubilize BaxWT and BaxP168A. The properties of the resolubilized proteins were not significantly different from the properties of His_6_-Bax directly synthesized in the presence of Brij-58 or F8-TAC. CD spectra showed that both proteins were organized as α-helices (~70%), corresponding to the measurements on His_6_-tagged proteins [28] and to the proportion that can be deduced from the structure (66%) [9]. Both proteins were able to bind to liposomes in a flotation assay, and to form pores, with a size large enough to permeabilize the membranes to 10kDa-dextran, which is close in size to cytochrome c (12.5kDa). Both proteins were also able to permeabilize yeast mitochondria to cytochrome c. Interestingly, in all these experiments, BaxP168A was, as expected, slightly more active than BaxWT, a property that has been previously observed for the His_6_-tagged proteins synthesized and maintained in solution in the presence of F8-TAC [28].

The next step was to reconstitute resolubilized Bax into nanodiscs. This was done by a classical detergent removal approach. A SEC separation allowed to identify fractions that were enriched with Bax-containing nanodiscs over fractions containing empty nanodiscs. Furthermore, we observed that BaxP168A-containing nanodiscs were slightly larger than BaxWT-containing nanodiscs, when observed by DLS. This was next confirmed by TEM observations showing a significant increase of the size of BaxP168A-containing nanodiscs compared to BaxWT-containing nanodiscs. Interestingly, the small amount of nanodiscs that were formed during the cell-free expression of His_6_-Bax in the presence of StrepII-MSP1 nanodiscs exhibited the same difference of size distribution between BaxWT and BaxP168A.

As reported previously, BaxP168A spontaneously forms pores in the OMM when it is expressed in yeast [27] or human cells [26]. The recombinant His_6_-tagged BaxP168A was slightly but significantly more efficient to form pores in liposomes and isolated mitochondria [28] and, in the present study, we show that the resolubilized proteins was also more active, both on liposomes and mitochondria. Then, for the first time, we can correlate this biochemical properties of BaxP168A to its capacity to form larger objects in membranes.

Although the size of nanodiscs is expected to be limited by the presence of the belt protein MSP1, they seem to have enough flexibility to adapt to the size of BaxP168A oligomers. It has been shown that the thickness of the lipid bilayer and the density of lipid molecules is not homogenous even in nanodiscs containing only lipids [57]. When a protein is inserted, the shape of nanodiscs may change. For example, nanodiscs having inserted bacteriorhodopsin adopt a squared shape [58]. SAXS measurements show that nanodiscs can also adopt elliptical shapes [59]. It is then likely that the size, and possibly the shape, of BaxP168A oligomers may create physical constraints forcing nanodiscs to adapt their size and shape.

One study has previously reported the reconstitution of Bax in nanodiscs [60]. His_6_-Bax was previously activated with a Bid-BH3 peptide and co-formed with DPPC/MSP1 nanodiscs and observed by cryo-EM. The size and the shape of Bax-containing nanodiscs were different from empty nanodiscs, with a more ellipsoidal shape and the presence of a pore having a diameter of 3.5nm. The authors assumed that Bax monomers were inserted [61]. However, the treatment of Bax by a Bid-BH3 peptide that was used in this study was later shown to induce the formation of, at least, a Bax dimer [34]. Molecular dynamics simulations of Bax inserted in nanodiscs have also been done [62]. From these data, the addition of a single Bax protein in the lipid bilayer would be sufficient to trigger the formation of a pore, by reorganizing lipid molecules in the nanodisc in close proximity to Bax, thus forming some empty space as a side effect. However, the conclusion that a monomer, or even a dimer, of Bax is sufficient to form a pore having a size sufficient to permeabilize a membrane to cytochrome c is not in line with electrophysiology experiments, in which the formation of Bax oligomers is needed to form the pore [6].

In our experiments, the difference of surface between BaxWT and BaxP168A nanodiscs is about 58nm^2^ (considering they are actual discs). The average diameter of an α-helix is about 1.2nm. Assuming a model where 3 helices are inserted (α5, α6 and α9) and Bax oligomers are formed of 12 monomers [15], the inserted helices would occupy a surface of about 40nm^2^. This would let a surface of about 18nm^2^ for the pore (about 4.8 nm in diameter), that is consistent with its expected size calculated from electrophysiology experiments: indeed, the Bax pore is permeable to cytochrome c (~3 nm in diameter), to ribonuclease A (~3.5 nm in diameter), but not to hemoglobin (~5 nm in diameter) [6]. Consistently, Guihard *et al* [35] calculated a diameter of 4.6nm for the Bax pore in liver mitochondria of rats developping fulminant hepatitis. This size is significantly larger than the 3.5nm pore depicted in Bid-BH3-activated Bax nanodiscs [60–62], that might correspond to an earlier step in the process of the pore formation, before Bax is fully oligomerized. It is noteworthy that a 3.5nm-pore would be theoretically large enough to permeabilize the membrane to cytochrome c [61], but would be very limiting to permeabilize the membrane to smac/diablo, that is released more slowly from mitochondria [37], and may require additional steps, compared to cytochrome c [63]. In the case of BaxP168A, which spontaneously forms oligomers [28], the initial steps corresponding to the insertion of a monomer/dimer might be absent, and a full size oligomeric pore would then be formed (Fig.7).

**Figure 7:**
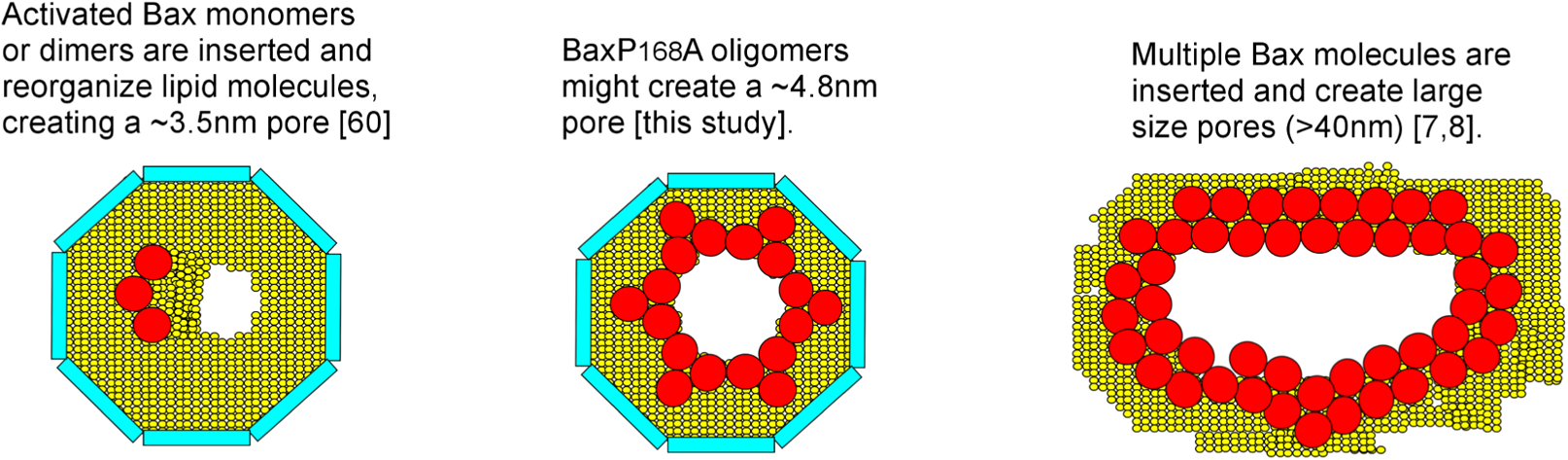
Schematic representation of different Bax pores. **Left:** Recombinant wild-type His6-Bax activated by the Bid-BH3 peptide forms ~3.5nm pores when co-formed in nanodiscs [60]. The reorganisation of lipid molecules in close proximity to Bax is sufficient to generate the pore [61]. **Right:** Endogenous Bax in actinomycin D-treated U2OS cells [7], or GFP-Bax expressed in HeLa cells [8] form large ring structures (40 to 100nm in diameter) in the OMM. Heat-activated recombinant Bax form similar large ring structures observed by AFM [8]. **Center:** Recombinant BaxP168A oligomers generate nanodiscs having a size compatible with the existence of a ~4.8nm pore, consistent with electrophysiology experiments [6,35].

Even though the BaxP168A pore would be larger than the Bid-BH3-activated Bax pore, it remains much smaller than the large objects forming rings having a diameter of several tenth of nanometers that were observed by high resolution fluorescence microscopy and atomic force microscopy [7,8]. These large objects might correspond to later apoptotic steps, when multiple Bax molecules would come together to form large, unselective holes in the OMM [36], while the “small” pores may correspond to the early steps of apoptosis, when the permeabilization is limited to several apoptogenic factors, namely cytochrome c and smac/diablo (Fig.7). The advantages of nanodiscs is that the size constraints imposed by the scaffold protein may limit the reconstitution, or the formation, of these large objects. On the basis of data reported in this paper, we will next optimize the method to obtain a higher number of observable objects allowing to determine a faithful 3D structure of BaxP168A oligomers reconstituted in nanodiscs.

## Experimental procedures

### Membrane Scaffold Protein (His_6_-MSP1E3D1) purification

The plasmid pET28b encoding His_6_-tagged MSP1E3D1 was transformed into *E.coli* BL21DE3* and plated on LB solid medium (0.5% Yeast Extract, 1% Tryptone, 1% NaCl, 1.5% Agar) containing 30mg/L kanamycin. One isolated colony was pre-grown in LB medium containing 30mg/L Kanamycin overnight at 37°C. The preculture was inoculated into 1L of TB medium (2.4% Yeast Extract, 1.2% Tryptone, 0.5% glycerol, 8.9mM Sodium Phosphate buffer pH 7.0) containing 30mg/L of Kanamycin at OD_600_ = 0.02 and allowed to grow until OD_600_= 0.9. MSP1E3D1 expression was induced by adding 1mM IPTG, and the growth was extended at 37°C until OD_600_ reached 1.8. Cells were pelleted by centrifugation (4,000 x g, 10 min, 4°C, Sorvall SLA1500 rotor), then resuspended in 30mL lysis buffer (100mM Na_2_HPO_4_, pH 7.6, 50mM NaCl, 1mM PMSF, 1mM Pic8340, 1mM AEBSF, 1% Triton X100, one tablet of Complete EDTA-free). Cells were lysed by sonication (50% pulsation, Power on 7 (max microTip), 7 cycles of 30 s of sonication followed by 30 s pause). Cells debris were pelleted by centrifugation (46,000 x g, 30 min, 4°C, Beckman 45Ti rotor). The supernatant was loaded overnight (in a closed circuit on a peristaltic pump) on a Ni-NTA column (HisTrap FF 5mL, Cytiva) previously equilibrated in lysis buffer. The circuit was opened and the column was washed with 5 volumes of buffers 1 to 3 (Buffer 1: 40mM Tris/HCl pH 8.0, 300mM NaCl, 1% Triton X100; Buffer 2: 40mM Tris/HCl pH 8.0; 300mM NaCl, 50mM Sodium Cholate; Buffer 3: 40mM Tris/HCl pH 8.0, 300mM NaCl, 50mM imidazole). The column was then coonected to an Äkta purifier and washed one more volume of buffer 3. Proteins were eluted with elution buffer (40mM Tris/HCl pH 8.0, 300mM NaCl, 300mM imidazole). The eluate was then dialyzed against the desalting buffer (10mM Tris/HCl pH 8.0, 100mM NaCl) and concentrated on Vivaspin concentrator (10kDa cut-off). The protein (~7mg/mL) was stored as working aliquots at −80°C.

A StrepII-tagged version of MSP1E3D1 was also purified, starting from a pET28b plasmid expressing StrepII-TEV-MSP1E3D1. The bacterial lysate was obtained as above and was loaded on a 1mL Strep-TactinXT-Superflow column (IBA) and connected to an Äkta Purifier. The column was washed with 5mL of 10mM Tris/HCl (pH 8.0) containing 150mM NaCl and 1mM EDTA, 5 mL of the same buffer added with 1% Triton X-100, and 5mL of the same buffer added with 50mM sodium cholate. The protein was then eluted with 5mL of the same buffer added with 50mM biotine. Fractions containing StrepII-TEV-MSP1E3D1 were identified after a SDS-PAGE revealed by Coomassie Blue staining and pooled. They were loaded on a 16/600 Superdex 75 column and Size Exclusion Chromatography (SEC) was done with a 20mM Tris/HCl buffer (pH 8.0) containing 100mM NaCl and 1mM EDTA. The fractions containing the protein were pooled and concentrated on Vivaspin concentrators (10kDa cut-off). The protein (~7mg/mL) was stored as working aliquots at −80°C.

### Nanodiscs preparation

2.5mg of 1-palmitoyl-2-oleoyl-sn-glycero-3-phosphocholine (POPC) in chloroform (Anatrace) were dried for 3h under argon flow. The dry lipid film was resolubilized in nanodisc buffer (10mM Tris/HCl, pH 8.0, 100mM NaCl, 200mM Sodium Cholate). 1.5mg of the Membrane Scaffold Protein (His6-MSP1E3D1) was added and left for equilibration for 1h. Detergent was removed by adding 150mg of BioBeads SM-2 (Bio-Rad) for 3h. The sample was then centrifuged at 20,000 x g for 15 minutes at 4°C, filtered through a 0.22µm membrane and loaded on a SEC column (Superdex 200 10/300GL, GE Healthcare). SEC was run in the same buffer (without cholate) at 0.5mL/min. When needed, nanodiscs were concentrated on Vivaspin with a 30kDa cut-off (final concentration: 0.2 to 0.4mM MSP1).

### Bax cell-free synthesis

Human Bax cDNA was cloned in a pIVEX 2.4 plasmid (Roche). The cloning was done so that the His_6_ tag was absent from the construction. Cell-free protein synthesis [64] was done in a small volume dialysis chamber (100 µL), separated from a feeding reservoir (1,700µL) by a dialysis membrane (MW 10,000). The system was routinely set up in an inverted microcentrifuge tube. Both chambers contain 0.1M K/Hepes (pH 8.0), 1mM EDTA, 0.05% NaN_3_, 2% PEG 8000, 151mM Potassium Acetate, 7.1mM Magnesium Acetate, 0.1mg/mL Folinic acid (calcium salt), 2mM DTT, 1mM (each) NTP mix, 0.5mM (each) aminoacid mix, 1mM (each) RDEWCM mix, 20mM K/PEP, 20mM acetylphosphate (Li/K salt), protease inhibitors cocktail (Complete without EDTA, Roche). The dialysis chamber was supplemented with components for the synthesis: 35% (v/v) S30 *E.coli* BL21DE3 lysate (in 10mM tris-acetate buffer, pH 8.2, 14mM Magnesium Acetate, 0.6mM Potassium Acetate, 0.5mM DTT), 0.04mg/mL Pyruvate Kinase (Sigma), 15µg/mL pIVEX-Bax plasmid, 0.5 mg/mL tRNAs mix (Roche), 6 units/mL T7 RNA polymerase [65], 3 units/mL RNasin (Promega). Depending on the experiments (see results), preformed nanodiscs were added. The synthesis was done for 16 hours at 30°C under gentle agitation.

The reaction mix was centrifuged for 15 min at 20,000 x g. In the presence of pre-formed nanodiscs, a large proportion of Bax protein was found in the pellet (see results). Bax was next resolubilized in the presence of Nonidet-P40 or Brij-58 (see results).

### Bax-containing nanodiscs co-formation

2.5mg of of a POPC/POPG 3/1 (w/w) mixture in chloroform (Anatrace) were dried for 3h under argon flow. The dry lipid film was resolubilized in co-formation buffer (20mM Tris/HCl, pH 8.0, 200mM NaCl, 200mM Sodium Cholate). BaxWT or BaxP168A resolubilized in Brij-58 (see results) were treated on detergent removal spin columns (Pierce) to remove the excess of Brij-58. The protein recovery yield was about 50%. The rest of the protocol was similar to empty nanodiscs (see above).

The insertion of His_6_-BaxWT and His_6_-BaxP168A into nanodiscs formed with StrepII-MSP1E3D1 was done by cell-free synthesis in the presence of pre-formed nanodiscs prepared with POPC and StrepII-MSP1E3D1 (see above). Like for the synthesis of untagged Bax in the presence of His6-MSP1E3D1, a large proportion of Bax was precipitated. Nevertheless, the supernatant containing nanodiscs (0.5mL) was mixed with StrepII-Tactin sepharose (IBA) and incubated overnight at 4°C under gentle agitation. Sepharose beads were washed several times with a 100mM Tris/HCl buffer (pH 8.0) containing 150mM NaCl and 1mM EDTA. The elution was done by adding the same buffer containing 50mM Biotin and incubated overnight at 4°C under gentle agitation. The supernatant was collected and dialysed against a 20mM Na/Hepes buffer (pH 7.8) containing 150mM NaCl. The dialysate was then added to Ni-NTA superflow resin (Qiagen) and incubated overnight at 4°C under gentle agitation. The resin was washed twice with the same buffer, and the elution was done with the same buffer containing 250mM imidazole. The eluate was dialyzed against the same buffer without imidazole, and concentrated with Vivaspin concentrators (cut-off 30kDa).

In all the experiments, Bax preparations were stored at 4°C and used rapidly (within one week).

### Recombinant cBid and Bcl-xL production

Recombinant human His_6_-Bid was produced in *E.coli* strain BL21DE3* transformed with the expression plasmid pET23d-His_6_-Bid, purified, and cleaved with caspase-8 as described by Kudla *et al* [66]. The protein was added with 15% glycerol and stored at −80°C in small aliquots.

Recombinant human His_6_-Bcl-xL was produced in *E.coli* strain BL21DE3* transformed with the expression plasmid pET28a-His_6_-Bcl-xL, as described by Yao *et al* [67]. The lysate contained folded His_6_-Bcl-xL deprived of the 15 C-terminal residues. The protein was added with 15% glycerol and stored at −80°C in small aliquots.

### LUV preparation process and dextran release assay

3mg of a phospholipid mixture mimicking mitochondrial lipid (46.5% POPC, 28.5% POPE, 9% DOPS, 7% Cardiolipin w/w/w/w) [68] was dried for 3h under argon flow. The dry lipid film was re-hydrated in a LUV buffer (25mM Na/Hepes pH 7.5, 200mM NaCl) supplemented with 10mM FITC-Dextran 10kDa (Sigma). The mixture was sonicated in a water bath for 3 minutes and extruded through a 400nm pore size Nucleopore Track-Etched membrane in a Liposofast extruder (Avestin). LUV were applied on a 10mL Sephadex-G50 gravity flow column. 1mL fractions were collected and the LUV containing fractions were identified by their cloudy appearance.

The Dextran release assay was based on a previously published protocol [69]. A mixture containing 20µL LUV and 8µg anti-FITC antibody (Thermofisher) was prepared. The assay was performed at 30°C using 96-well plates. 100ng of resolubilized BaxWT or BaxP168A were passed through a detregent-removal spin column and added to liposomes (100μL final volume). FITC fluorescence was measured every 1 min during 60–90min in a microplate fluorescence reader (Clariostar, BMG Labtech). The excitation and emission wavelengths were set at 495nm and 521nm, respectively. The 0% of fluorescence decay was measured on the mixture of LUV and anti-FITC antibody alone, and the 100% of decay was measured after adding 0.5% CHAPS.

### Flotation assay

LUV were prepared as above, without FITC-dextran. 30% and 80% (w/w) Histodenz (Sigma) stock solutions were prepared in LUV buffer. 100 µL of resolubilized BaxWT (~ 5 µg) were mixed with 100 µL of LUV, completed to 1mL with LUV buffer, and incubated for 1-2 hours at 4°C. 750 µL of the 80% Histodenz solution were added and thoroughly mixed with 750µL of the protein-LUV sample in Thinwall centrifuge tubes (Beckman 50.2ti rotor). 1.5 mL of the 30% Histodenz solution was added as the intermediate layer, and 1.5mL LUV buffer was added as the top layer. Tubes were centrifuged overnight at 30,000 x g at 4°C in a 50.2ti swinging bucket rotor in a Beckman L-60 ultracentrifuge. The gradients were fractionated in 500µL aliquots, precipitated with 0.3M Trichloroacetic acid, washed with acetone, dried, resolubilized in Laemmli buffer, and analyzed by western blotting against Bax.

### Circular Dichroism (CD)

Samples for CD measurement were dialyzed against 40mM Sodium Phosphate buffer (pH7.2) in dialysis cassettes (Slide-A-Lyzer, Thermofisher) to remove Cl^−^ ions. Far UV spectra (190-260 nm) were measured on a Jasco J-810 CD spectropolarimeter. The acquisition was carried out at 25°C with a 1mm optical path length cell under nitrogen atmosphere. The fractions obtained after the dialysis were diluted to 65-100 µg/mL in 200µl. 10 scans were collected from 260 to 190 nm at 0.5 nm intervals, then averaged and baseline-corrected by substracting buffer spectra. They were analyzed at Dichroweb (http://dichroweb.cryst.bbk.ac.uk) using the CDSSTR algorithm [70] with the SP175 dataset as a reference [71].

### Dynamic Light Scattering (DLS)

The DLS of eluted nanodisc-containing fractions from SEC was measured in a Wyatt Dynapro Nanostar, using the standard mode of acquisition, in Eppendorf UVettes. The time delay curves were examined and the boundaries were slightly adjusted if needed.

### Mitochondria permeabilization

Mitochondria from the yeast wild-type strain W303-1B were isolated and Bax permeabilization assay were done as described previously [28]. Briefly, mitochondria were suspended at 1mg/mL in 0.1mL of PMB buffer (10mM K/Hepes pH 7.4, 250mM sucrose, 80mM KCl, 2mM MgOAc, 1mM KH_2_PO_4_, 1mM ATP, 0.08mM ADP, 5mM succinate). After Bax and adequate controls were added, samples were incubated for 1 hour at 30°C. Samples were then centrifuged (25,000 x g, 10 min, 4°C). The mitochondrial pellet was resuspended in 100µL water and mitochondrial proteins were precipitated with 0.3M Trichloroacetic acid. The pellet was washed with acetone, dried and resolubilized in Laemmli buffer.

### Immunoprecipitation and co-immunoprecipitation

For immunoprecipitation assays, 50ng of Bax-containing nanodiscs were added with 1µg of anti-Bax 2D2 or anti-Bax 6A7 mouse monoclonal antibodies (Santa-Cruz) and incubated overnight at 4°C. 50µL of a suspension of Protein G coupled to agarose beads (Sigma) previously equilibrated in Bax buffer (20mM Tris/HCl pH 8.0; 200mM NaCl) were added and incubated for 3h. Agarose beads were washed 4 times on IP columns (Sigma) with a washing buffer (25mM Tris/HCl pH 7.4, 500mM NaCl, 1mM EDTA, 1% Nonidet P40, 10% glycerol) and twice with the same buffer at a 0.1X dilution. Bound proteins were recovered by adding a small volume of Laemmli buffer without β-mercaptoethanol and a 15 min incubation at 65°C.

For co-immunoprecipitation assays, 50ng of Bax-containing nanodiscs were incubated with 100ng of Bcl-xL for 1h. 1µg of mouse monoclonal anti-Bax antibody (2D2, Santa-Cruz) or rabbit monoclonal anti-Bcl-xL antibody (E18, Abcam) were added and incubated overnight at 4°C. The remaining steps were as above.

### Transmission Electron Microscopy (TEM)

Nanodiscs were negatively stained according to Daury *et al* [72]. Briefly, sample was applied to a glow-discharged carbon-coated copper 300 mesh grids and stained with 2% uranyl acetate (w/v) solution. Images were recorded under low-dose conditions on a Tecnai F20 FEG electron microscope (FEI, ThermoFisher Scientific) operated at 200 kV using an Eagle 4k_4k camera (ThermoFisher Scientific).

The area of nanodiscs was measured manually with Image J (https://imagej.nih.gov/ij/) and the dm3 reader plugin (https://imagej.nih.gov/ij/plugins/DM3_Reader.html). The average diameter was calculated by considering that they were perfect disks.

### Protein dosage, SDS-PAGE and Western-Blots

Protein concentrations was determined by the bicinchonic acid (BCA)-assay (Thermofisher). Protein samples were solubilized in Laemmli buffer (0.1M Tris/HCl pH 8.0, 2% SDS, 2% β-mercaptoethanol, 10% glycerol, 0.004% Bromophenol Blue) and heated at 65°C for 15 minutes. Samples were run on Tris/glycine 12.5% SDS PAGE and transferred onto 0.2 μm PVDF membranes (Hybond, Amersham). Membranes were saturated with 5% milk in PBS-Tween 20 (or TBS-Tween 20 for the anti-Bcl-xL antibody). Primary antibodies were added overnight at 4°C, and secondary antibodies were added for 45 min at room temperature. Antibodies were as follows: rabbit polyclonal anti-human Bax N20 (Santa-Cruz) 1/5,000 dilution, mouse monoclonal anti-human Bax 2D2 (Santa-Cruz) 1/5,000 dilution, rabbit monoclonal anti-Bcl-xL antibody (E18, Abcam) 1/10,000 dilution, mouse monoclonal anti-his_6_ tag (Thermofisher) 1/10,000 dilution, mouse monoclonal anti-yeast porin (Thermofisher) 1/50,000 dilution, rabbit polyclonal anti-yeast cytochrome c (custom antibody, Millegen) 1/5,000 dilution, peroxidase-coupled goat anti-rabbit IgG (Jackson Immunoresearch) 1/10,000 dilution, peroxidase coupled goat anti-mouse IgG (Jackson Immunoresearch) 1/10,000 dilution. Peroxidase activity was revealed by Enhanced Chemiluminescence (Luminata Forte, Millipore), recorded with a digital camera (G-Box, Syngene) and analyzed with Image J (https://imagej.nih.gov/ij/). Coomassie Blue staining was done by soaking the gels in ethanol/water/acetic acid 5/5/1 (v/v/v/) for 30 minutes, staining with 0.25% Coomassie Brillant Blue G250 (Sigma) in the same mixture for 30 minutes, and destaining in the same mixture until the signal to background ratio is satisfactory. Plots were done with Graphpad Prism 6.05 software with the integrated statistical analysis tools.

## Aknowledgements

The work was supported by the Centre National de la Recherche Scientifique (UMR5095 and UMR5248), the Université de Bordeaux (UMR5095 and UMR 5248) and the Ligue Régionale contre le Cancer (to S.M. and C.G.). A.R.E. is recipient of a PhD fellowship from the Agence Nationale des Bourses du Gabon.

The authors declare that they have no conflicts of interest with the contents of this article.

## Abbreviations

AEBSF: 4-(2-Aminoethyl)benzenesulfonyl fluoride
ANTS: 8-Aminonaphthalene-1,3,6-trisulfonate
CN-PAGE: Colorless Native PAGE
DLS: Dynamic Light Scattering
DPX: p-Xylene-bis(N-pyridinium bromide)
IPTG: Isopropyl β-D-thiogalactoside
LB: Luria-Bertani medium
MLV: Multilamelar Vesiccles
MSP1: Membrane Scaffold Protein
OMM: Outer Mitochondrial Membrane
PMSF: phenylmethylsulfonyl fluoride
POPC: 1-palmitoyl-2-oleoyl-sn-glycero-3-phosphocholine
POPG: 1-palmitoyl-2-oleoyl-sn-glycero-3-phosphoglycerol
SEC: Size Exclusion Chromatography
TB: Terrific Broth
TEM: Transmission Electron Microscopy

